# Conserved RGF1 peptide signaling regulates root meristem development through ROS in *Arabidopsis* and rice

**DOI:** 10.64898/2026.07.06.736907

**Authors:** Joon-Keat Lai, Jhen-Ni Jhang, Hong-Chun Yen, Hsin-Yen Cho, Yu-Chun Hsiao, Hariharan Balasubramaniam, Ching-Shan Tseng, Masashi Yamada

## Abstract

The root meristem is essential for stem cell maintenance and root development in plants. In *Arabidopsis*, Root meristem Growth Factor (RGF) peptides and their receptors regulate root meristem size through reactive oxygen species (ROS)-dependent signalling. RGF1-mediated ROS redistribution post-translationally stabilises the root meristem master regulator PLETHORA2 (PLT2). Although genomic studies suggest that RGF–receptor modules are evolutionarily conserved across land plants, their functional characterisation has remained largely limited to *Arabidopsis*. Here, we show that *Oryza sativa* RGF1-1 (OsRGF1-1) functions as a rice homologue of *Arabidopsis* RGF1 (AtRGF1). CRISPR/Cas9-generated *Osrgf1-1* mutants exhibited shorter seminal roots, reduced root meristem size, and decreased superoxide (O₂•⁻) accumulation. EdU staining further confirmed that cell proliferation activity was reduced in the *Osrgf1-1* mutants. The *Osrgf1-1* mutants were sensitive to low concentrations of chemically synthesised mature OsRGF1-1 peptide. This low dose of OsRGF1-1 peptide restored seminal root growth and O₂•⁻ accumulation in the *Osrgf1-1* mutants but had no detectable effect on the wild type. Functional analyses using *Arabidopsis rgfr* receptor mutants further demonstrated that OsRGF1-1 is perceived through conserved RGF receptor machinery. Together, our findings provide the first functional evidence that the RGF1–receptor–ROS signalling module is evolutionarily conserved between dicots and monocots in the regulation of root meristem development.

## Introduction

The root meristem continuously generates new cells while maintaining stem cell activity throughout the plant’s life. In *Arabidopsis*, root meristem development is regulated by the Root meristem Growth Factor (RGF; also known as GOLVEN or CLE-like) peptide signaling pathway (Matsuzaki *et al*., 2010; Meng *et al*., 2012; Whitford *et al*., 2012), which controls meristem size through reactive oxygen species (ROS)-dependent mechanisms (Yamada *et al*., 2020). The *RGF1/2/3* genes are expressed around the root apical meristem and function redundantly in root meristem development (Matsuzaki *et al*., 2010). Although single *rgf* mutants do not exhibit obvious root phenotypes, the *rgf1/2/3* triple mutant forms a smaller root meristem, which can be rescued by treatment with a synthetic sulfated RGF1 peptide. Receptors for RGF1 have been identified and localised to the meristematic zone (Ou *et al*., 2016; Shinohara *et al*., 2016; Song *et al*., 2016). Mutants defective in RGF receptors (also known as RGF1 INSENSITIVE) are less sensitive to RGF1 peptide treatment, demonstrating that the RGF–receptor pathway maintains and modulates root meristem size (Ou *et al*., 2016; Shinohara *et al*., 2016; Song *et al*., 2016).

The spatial distribution of ROS across distinct root developmental zones plays an important role in root development. Superoxide (O₂•⁻) and hydrogen peroxide (H₂O₂) accumulate predominantly in the meristematic zone and elongation/differentiation zone, respectively, where they regulate developmental progression (Dunand *et al*., 2007; Tsukagoshi *et al*., 2010). The balance and spatial distribution of these ROS are dynamically modulated by developmental and environmental cues. The RGF1–receptor pathway regulates the distribution of O₂•⁻ and H₂O₂ in the root meristem (Yamada *et al*., 2020). Altered ROS distribution downstream of RGF1 signalling post-translationally regulates the stability of PLETHORA2 (PLT2), a master regulator of root meristem activity (Hsiao *et al*., 2025; Yamada *et al*., 2020).

Although root meristems across land plants share common developmental functions, their anatomical organisation differs substantially between species. Rice roots possess multiple cortical cell layers and exhibit a stem-cell niche architecture distinct from that of *Arabidopsis*, reflecting major evolutionary divergence between monocot and dicot root systems (Rebouillat *et al*., 2009; Wang *et al*., 2014). Despite these structural differences, comparative genomic analyses have suggested that the RGF–receptor signalling module is conserved across diverse land plant lineages (Fang *et al*., 2021; Ghorbani *et al*., 2015). However, functional evidence supporting the conservation of RGF signalling outside *Arabidopsis* remains limited. Whether RGF-mediated control of root meristem development and ROS homeostasis represent a conserved regulatory mechanism across evolutionarily divergent plant species therefore remains unclear.

Previous comparative genomic and phylogenetic analyses predicted the presence of RGF1 homologues in the *Oryza sativa* genome (Fang *et al*., 2021; Ghorbani *et al*., 2015). However, the molecular identity and biological functions of *Oryza sativa* RGF1 peptides have remained uncharacterised. Here, we identify OsRGF1-1 (LOC_Os02g09760) as a functional rice homologue of *Arabidopsis* RGF1. Genetic, physiological, and peptide application analyses demonstrate that OsRGF1-1 regulates root meristem size and ROS distribution through a conserved receptor-dependent signalling pathway in rice. These findings demonstrate that RGF-mediated ROS signalling is evolutionarily conserved between *Arabidopsis* and rice and establish OsRGF1-1 as a regulator of root meristem development in a monocot species with a root meristem architecture distinct from that of *Arabidopsis*.

## Materials and Methods

### Plant material and rice growth conditions

All mutants and transgenic lines used in this study were in the *Oryza sativa* L. ssp. japonica variety Nipponbare background. Seeds were dehusked and heat-treated at 45°C for 24 h before use. Seeds were surface-sterilised in 50% bleach coupled with 0.1% Tween 20 (Sigma-Aldrich, USA) for 20 min, then rinsed ten times with sterile water. The sterilised seeds were then imbibed in sterile water for 8 h followed by imbibition in 1% (v/v) Plant Preservative Mixture (PPM™) (Plant Cell Technology, USA) for 16 h. The seeds were sown on half-strength Murashige and Skoog (MS) medium (PhytoTech Labs, USA) supplemented with 0.05% (v/v) PPM™ and 0.05% (w/v) MES hydrate (Sigma-Aldrich, USA). The medium pH was adjusted to 5.7 using 5 M potassium hydroxide (Sigma-Aldrich, USA). The medium was solidified by 0.15% (w/v) Gelzan™ agar (PhytoTech Labs, USA). The sown seeds were incubated in the dark for 1 day, followed by growth under 700 µmol m⁻² s⁻¹ light for 4 days at 28/22°C (day/night) in a growth chamber.

### Plant material and *Arabidopsis* growth conditions

The *rgfr123 Arabidopsis thaliana* (L.) Heynh. mutant used in this study is in the Columbia-0 (Col-0) genetic background and has been previously characterised (Shinohara *et al*., 2016). Seeds were surface-sterilised in 50% bleach supplemented with 0.1% Tween 20 (Sigma-Aldrich, USA) for 15 min, followed by five rinses with sterile distilled water. The sterilised seeds were then stratified at 4°C in the dark for 2 days. Subsequently, seeds were sown on half-strength Murashige and Skoog (½ MS) medium (Caisson Laboratories, USA) supplemented with 0.05% MES, 1% sucrose, and 1% agar (Sigma-Aldrich, USA), and the pH was adjusted to 5.7 with KOH. Seedlings were grown vertically in a growth chamber at 22°C under long-day conditions (16 h light/8 h dark) for 7 days. After this period, seedlings were transferred to ½ MS agar plates containing either sterile water (mock treatment), synthetic sulfated AtRGF1 peptide, or synthetic sulfated OsRGF1-1 peptide (Mission Biotech, Taiwan) for 24 h.

### Light and confocal microscopy analysis for *Arabidopsis* roots

For superoxide (O₂•⁻) detection, seedlings after mock or peptide treatments were stained with 400 μM nitroblue tetrazolium (NBT) dissolved in 20 mM phosphate buffer (pH 6.1) in the dark for 2 min, followed by two rinses with distilled water. Root images were captured using a 10× objective lens on a Zeiss AXIO Scope A1 microscope. Total NBT signal intensity within the root tip region was quantified using Fiji ImageJ and used for statistical analysis.

Root meristems were observed 24 h after mock or peptide treatment using a 20× objective on a Zeiss LSM 980 laser scanning confocal microscope. Prior to imaging, roots were stained with propidium iodide (PI) to visualise cell boundaries, with excitation at 561 nm and emission detected at 565–747 nm. Confocal images were assembled from multiple stitched tiles using a semiautomated mosaicking approach (Thevenaz and Unser, 2007). The meristematic zone was defined as the region extending from the quiescent centre (QC) to the first cortex cell whose longitudinal length exceeded twice that of the preceding cell, and the number of cortex cells within this region was counted to estimate meristem size.

### Generation of CRISPR/Cas9 *Osrgf1-1* mutants

Three single guide RNAs (sgRNAs) targeting the coding region of OsRGF1-1 were designed for CRISPR/Cas9-mediated mutagenesis. The target sequences were 5′-TGTCAGTGTAACCAGAGCAA-3′, 5′-TGAGGAAATGCAAGAACGGG-3′, and 5′-GAGGATGCCGACGAACTGAG-3′. The oligonucleotide sequences used for sgRNA construction are provided in Supplementary Data 2, Table S2. The CRISPR/Cas9 expression vector pYLCRISPR/Cas9Pubi-H (Ma *et al*., 2015) was transformed into Agrobacterium tumefaciens strain EHA105 and subsequently introduced into calli derived from the japonica rice cultivar Nipponbare (wild type). Two independent homozygous Osrgf1-1 mutant lines (T3 generation, free of the CRISPR/Cas9 transgene) were obtained. The mutations were confirmed by Sanger sequencing of PCR products amplified using primer pairs flanking the CRISPR target region. The primer sequences used for genotyping are provided in Supplementary Data 2, Table S5.

### Root growth assay

For measuring seminal root length, entire seminal roots were cut and scanned with a scanner, followed by measurement using ImageJ. For detecting O₂•⁻, seminal roots were stained with 1 mg/mL NBT (Invitrogen, USA) dissolved in 10 mM phosphate buffer (pH 7.8) containing 10 mM sodium azide (Sigma-Aldrich, USA). A distinct NBT staining protocol was used for rice roots, as the higher tissue density of rice roots requires modified infiltration conditions compared with *Arabidopsis*. The NBT solution was pre-warmed to 30°C before use. Rice seedlings were suspended on Falcon tubes using self-made funnel-shaped filter paper, with the root tips immersed in the NBT solution. The staining process was performed under vacuum infiltration for 25 min at room temperature. The roots were rinsed five times with distilled water to terminate the NBT reaction. Root images were captured using a dissection microscope (Leica M205 FCA) with a digital camera (Leica DFC450 C). The root tip region used for quantification was defined based on NBT staining intensity thresholds using an in-house software. The NBT total intensity within this region was automatically quantified by the software. The software is available upon request.

### Expression analysis of *OsRGF1-1* in rice root zones by RT-qPCR

To perform RT-qPCR, roots were harvested from 6–7-day-old wild-type rice seedlings germinated on ½ MS medium. The spatial pattern of ROS accumulation along the root axis was first confirmed in a separate set of roots by NBT staining, which revealed intense superoxide accumulation at the root tip and the adjacent region, with the signal gradually decreasing away from the root apex. Because the exact cellular boundary of the meristematic zone is difficult to determine in rice roots due to their multilayered cortical structure, the NBT staining pattern was used as a guide for root dissection. A similarly sized root tip segment extending from the root tip to the region where NBT staining markedly decreased was precisely excised on a 2% agar plate containing ½ MS medium using an ophthalmic scalpel (Feather) under a dissection microscope (Leica M205 FCA). This segment was operationally designated as the meristematic zone (MZ) for both seminal and crown roots. The adjacent region, in which NBT staining was largely absent, was collected separately and designated as the differentiation zone (DZ). Approximately 20 root segments per fraction were pooled for each biological replicate, immediately snap-frozen in liquid nitrogen, and stored at −80°C.

Tissues were homogenized in RLT buffer supplied with the RNeasy Micro Kit (Qiagen, Germany), and total RNA was isolated according to the manufacturer’s instructions. Following DNase I treatment, cDNA was synthesized from 100 ng of total RNA using SuperScript IV Reverse Transcriptase (Invitrogen, USA). RT-qPCR was performed using a CFX Connect Real-Time PCR Detection System (Bio-Rad, USA) with SYBR-based detection. The primer sequences used for RT-qPCR are provided in Supplementary Data 2, Table S3. Expression levels of *OsRGF1-1* were normalized to *OsPP2AA*. Three biological replicates and three technical replicates were analyzed for each experiment, and the data are presented as the mean ± standard error (SE) of three biological replicates. Statistical differences between the MZ and DZ were evaluated using Student’s t-test (\*\**P* < 0.001; *P* < 0.05).

### Construction of the *OsRGF1-1::GUS* reporter vector

A 1,585 bp genomic fragment upstream of the *OsRGF1-1* start codon was amplified using the primers OsRGF1-1pro_F and OsRGF1-1pro_R. The primer sequences are provided in Supplementary Data 2, Table S4. The amplified fragment was cloned into the pENTR/D/TOPO vector (Invitrogen, USA). The resulting entry clone was recombined into the destination vector pMDC162 (Curtis and Grossniklaus, 2003) using the Gateway™ LR Clonase™ II Enzyme Mix (Invitrogen, USA) to generate the *OsRGF1-1::GUS* reporter construct.

### EdU staining

For meristematic zone size measurements, roots were incubated in 10 µM 5-ethynyl-2’-deoxyuridine (EdU; Invitrogen, USA) for 1 h under the same growth conditions. Root tips were subsequently fixed in 4% formaldehyde prepared in 1× phosphate-buffered saline (PBS; pH 7.4) containing 0.1% Triton™ X-100 (Sigma, USA) for 30 mins at room temperature. Fixed root tips were washed three times with 1× PBS for 10 mins each. EdU labeling was performed using the Click-iT® reaction cocktail (Invitrogen, USA) according to the manufacturer’s instructions. Images were acquired using a confocal microscope with a 10× objective at excitation/emission wavelengths of 488 nm/499–539 nm, 10% laser intensity, 700 V master gain, 1.0 digital gain, and 4 µm Z-stack intervals. Maximum-intensity projections were generated from the Z-stack images, and total EdU fluorescence intensity was quantified as an indicator of meristematic zone size.

### Statistical analysis and reproducibility

Experiments were independently repeated three times with similar results. No power analysis was performed to estimate the sample size. All statistical analyses were performed using R version 4.5.2 (http://www.r-project.org/). For two-sample comparisons, normally distributed data were analysed with a two-tailed *t*-test, and non-normal data were analysed with a Wilcoxon test. As an omnibus test, Welch’s analysis of variance (ANOVA) or an aligned rank transform ANOVA (ART) was performed, depending on data normality. As a post hoc test, the ART-Contrast test was applied to non-normal data, and the Games–Howell test, REGWQ test, or Tukey–Kramer test was performed on normal data, depending on data homoscedasticity and experimental design. In the box plots, circles represent individual data points; the horizontal line within boxes represents the median; the upper and lower hinges indicate the 75th and 25th percentiles (interquartile range [IQR]), respectively; and the whiskers show 1.5 × IQR extensions. The numbers below the boxes represent the sample sizes. Lowercase letters above the boxes indicate the compact letter display for pairwise comparisons, in which different letters indicate significant differences (*P* < 0.05) among groups. Asterisks above the boxes indicate the significance of two-sample comparisons (*, *P* < 0.05; **, *P* < 0.01; ***, *P* < 0.001; ns, non-significant [*P* > 0.05]).

## Results

### LOC_Os02g09760 (*OsRGF1-1*) is identified as the most likely rice homologue of *AtRGF1*

The mature AtRGF1/2/3 peptides are located in the C-terminal region of their precursor polypeptides and consist of 13 amino acids (Fig. 1A) (Matsuzaki *et al*., 2010). LOC_Os02g09760 (hereafter *OsRGF1-1*) and LOC_Os09g0474600 (*OsRGF1-2*) were previously predicted as RGF homologues in *Oryza sativa* based on comparative genomic and phylogenetic analyses (Ghorbani *et al*., 2015). The predicted mature OsRGF1-1 peptide consists of 13 amino acids and contains the conserved aspartic acid–tyrosine (DY) motif required for RGF activity, as well as the conserved HPP and N residues (Fig. 1A). These residues are conserved among previously identified AtRGF peptides (Fernandez *et al*., 2013; Matsuzaki *et al*., 2010) and are essential for tyrosine sulfation by *TYROSYLPROTEIN SULFOTRANSFERASE 1* (*TPST1*), which is required for RGF biological activity (Komori *et al*., 2009).

**Fig. 1.**
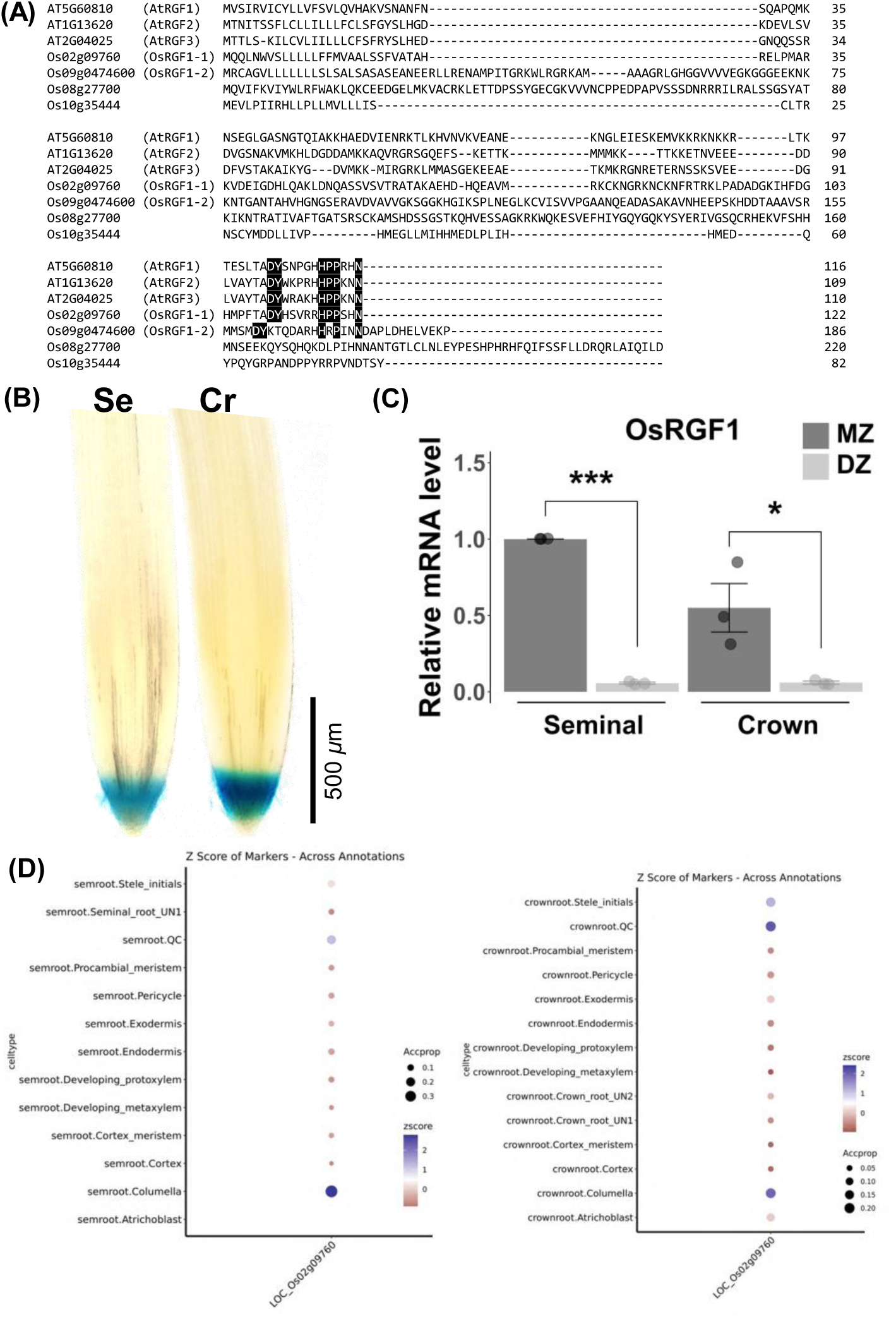
*OsRGF1-1* encodes a conserved RGF peptide expressed in the root meristematic zone. (A) Multiple sequence alignment of *Arabidopsis* RGF1 (AT5G60810), RGF2 (AT1G13620), RGF3 (AT2G04025), and *Oryza sativa* LOC_Os02g09760 (OsRGF1-1), LOC_Os09g0474600 (OsRGF1-2), LOC_Os08g27700, and LOC_Os10g35444. Conserved DY and HPP motifs are highlighted. (B) Histochemical GUS staining of transgenic rice seedlings expressing *pOsRGF1-1::GUS*. GUS activity is detected in the root tip of seminal (Se) and crown (Cr) roots. Bar = 500 µm (C) Relative OsRGF1-1 mRNA levels in the meristematic zone (MZ) and differentiation zone (DZ) of seminal and crown roots, determined by RT-qPCR. Data are shown as mean ± SE (n = 3 biological replicates). Asterisks indicate statistically significant differences between MZ and DZ (\*\*\**P* < 0.001; \**P* < 0.05, Student’s t-test). (D) Dot plots showing the Z-score of *OsRGF1-1* expression across cell type annotations in seminal and crown root single-cell RNA-seq datasets, analysed using the RiceSCBase platform. Dot size indicates the proportion of cells expressing OsRGF1-1; colour intensity indicates the mean Z-score.

In contrast, the predicted OsRGF1-2 mature peptide is not located in the C-terminal region and contains 15 amino acids rather than the canonical 13 amino acids. Given that mature RGF peptide length is highly constrained and that functionally conserved peptides typically retain both peptide length and key amino acid residues, OsRGF1-2 is unlikely to represent a canonical functional homologue of AtRGF1. Consistent with this interpretation, a later phylogenetic analysis placed OsRGF1-2 in a distinct clade separate from AtRGF1 and OsRGF1-1 (Fang *et al*., 2021), suggesting functional divergence. Although LOC_Os08g27700 and LOC_Os10g35444 were identified within the same clade as *AtRGF1/2/3* (Fang et al., 2021), both lack the conserved residues required for canonical RGF activity and are therefore unlikely to function as canonical RGF peptides (Fig. 1A). Together, these analyses indicate that OsRGF1-1 is the most likely rice homologue of AtRGF1. Because *AtRGF1/2/3* are specifically expressed around the *Arabidopsis* root apical meristem (Matsuzaki *et al*., 2010), we hypothesised that the putative rice homologue *OsRGF1-1* would likewise be expressed in the rice root meristem.

To examine the expression pattern of *OsRGF1-1*, we generated transgenic rice expressing a *β-glucuronidase* (*GUS*) reporter driven by the *OsRGF1-1* promoter (*pOsRGF1-1::GUS*) in Nipponbare (wild-type) plants. GUS signals were predominantly detected in the root apical regions of both seminal and crown roots (Fig. 1B). Reverse transcription quantitative PCR (RT-qPCR) analysis further confirmed that *OsRGF1-1* expression is preferentially enriched in the root apical meristem region compared with the differentiation zone (Fig. 1C). Publicly available rice single-cell RNA-seq datasets were analysed using the RiceSCBase platform (Yan *et al*., 2025; Zhang *et al*., 2021). Analysis of these datasets further supported the expression pattern observed by GUS staining and RT-qPCR, showing that *OsRGF1-1* is preferentially expressed in root meristematic cell populations (Fig. 1D). Collectively, these data indicate that *OsRGF1-1* is preferentially expressed in the root apical meristem region and represents the most likely rice homologue of AtRGF1. To investigate the biological function of OsRGF1-1 in root development, we generated loss-of-function mutants using CRISPR/Cas9.

### Generation of CRISPR/Cas9 mutants of *OsRGF1-1*

Three single guide RNAs (sgRNAs) were designed to target distinct regions of the third exon of *OsRGF1-1*, including the conserved region encoding the predicted mature peptide (Fig. 2A). PCR analysis of the *OsRGF1-1* locus showed that two independent mutant lines produced shorter amplicons than the wild type, consistent with CRISPR/Cas9-induced deletions (Fig. 2B). Sequencing revealed that *Osrgf1-1-7* carried 1-bp insertions at target sites 1, 2, and 3, together with a 26-bp deletion at target site 2 (Fig. 2C). These combined mutations introduced a premature stop codon upstream of the predicted mature peptide region, making production of the canonical mature OsRGF1-1 peptide highly unlikely (Fig. S1). The second allele, *Osrgf1-1-8*, contained a 55-bp deletion between target sites 1 and 2 and a 1-bp insertion at target site 3 (Fig. 2D). The 55-bp deletion caused a frameshift, removing 18 amino acids and altering both the putative processing region and the predicted mature peptide sequence (Fig. S1). As a result, the conserved aspartic acid–tyrosine (DY) motif required for RGF activity was eliminated (Fig. S1B). The additional 1-bp insertion at target site 3 further altered the downstream coding sequence.

**Fig. 2.**
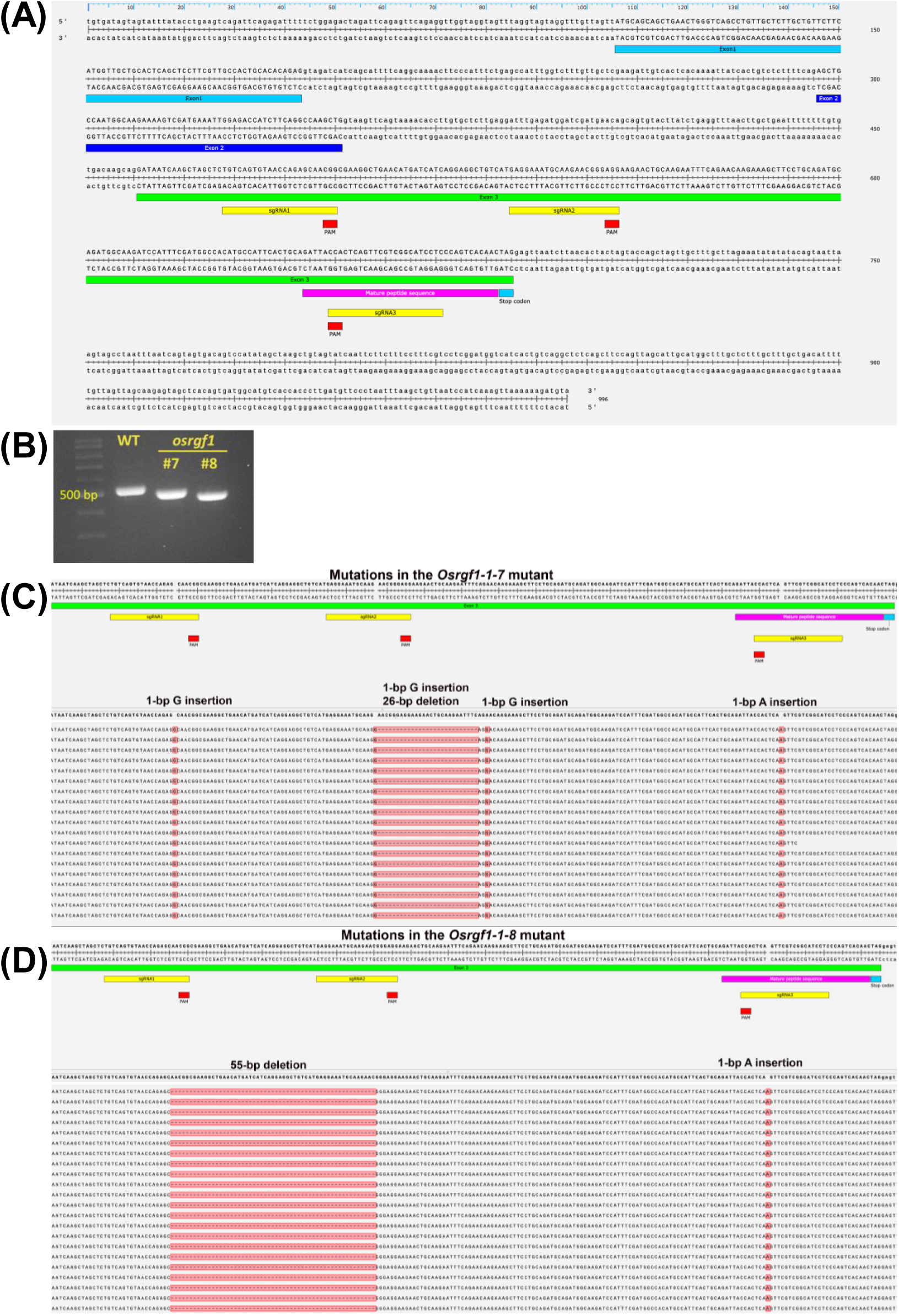
Molecular characterisation of *Osrgf1-1* mutant alleles. (A) Schematic of the *OsRGF1-1* genomic locus showing the positions of sgRNA1, sgRNA2, and sgRNA3 target sites and the mature peptide sequence. Exons are indicated. (B) PCR genotyping of wild-type (WT) and *Osrgf1-1* (#7 and #8) plants. (C) Sequencing alignment of the *Osrgf1-1-7* mutant showing the identified mutations: 1-bp G insertions at three sites, a 1-bp A insertion, and a 26-bp deletion. (D) Sequencing alignment of the *Osrgf1-1-8* mutant showing a 55-bp deletion and a 1-bp A insertion.

Thus, both mutant alleles are highly unlikely to produce a functional mature OsRGF1-1 peptide (Fig. S1).

### *Osrgf1-1-7* and *Osrgf1-1-8* exhibit defects in root growth, meristem activity, and O₂•⁻ accumulation

To characterise the root phenotypes of the *Osrgf1-1* mutants, we measured seminal root length in Nipponbare (wild type), *Osrgf1-1-7*, and *Osrgf1-1-8* at 8 days after transplantation (DAT). Both mutants produced significantly shorter seminal roots than the wild type (Fig. 3A, D). Because *OsRGF1-1* is expressed in the root meristem region (Fig. 1B, C, and D), we next examined whether loss of OsRGF1-1 affects root meristem activity. We used the extent of the EdU-positive proliferation domain to measure root meristem activity. Proliferating cells in seminal roots were detected using EdU (5-ethynyl-2′-deoxyuridine) staining. Both mutants showed weaker and more restricted EdU signals than the wild type, indicating a reduced number of proliferating cells and a smaller proliferation domain (Fig. 3B, E). Consistent with these observations, the *Atrgf1/2/3* triple mutant also exhibits reduced root meristem size (Matsuzaki et al., 2010), suggesting that OsRGF1-1 functions similarly to AtRGF1 in regulating root meristem development.

**Fig. 3.**
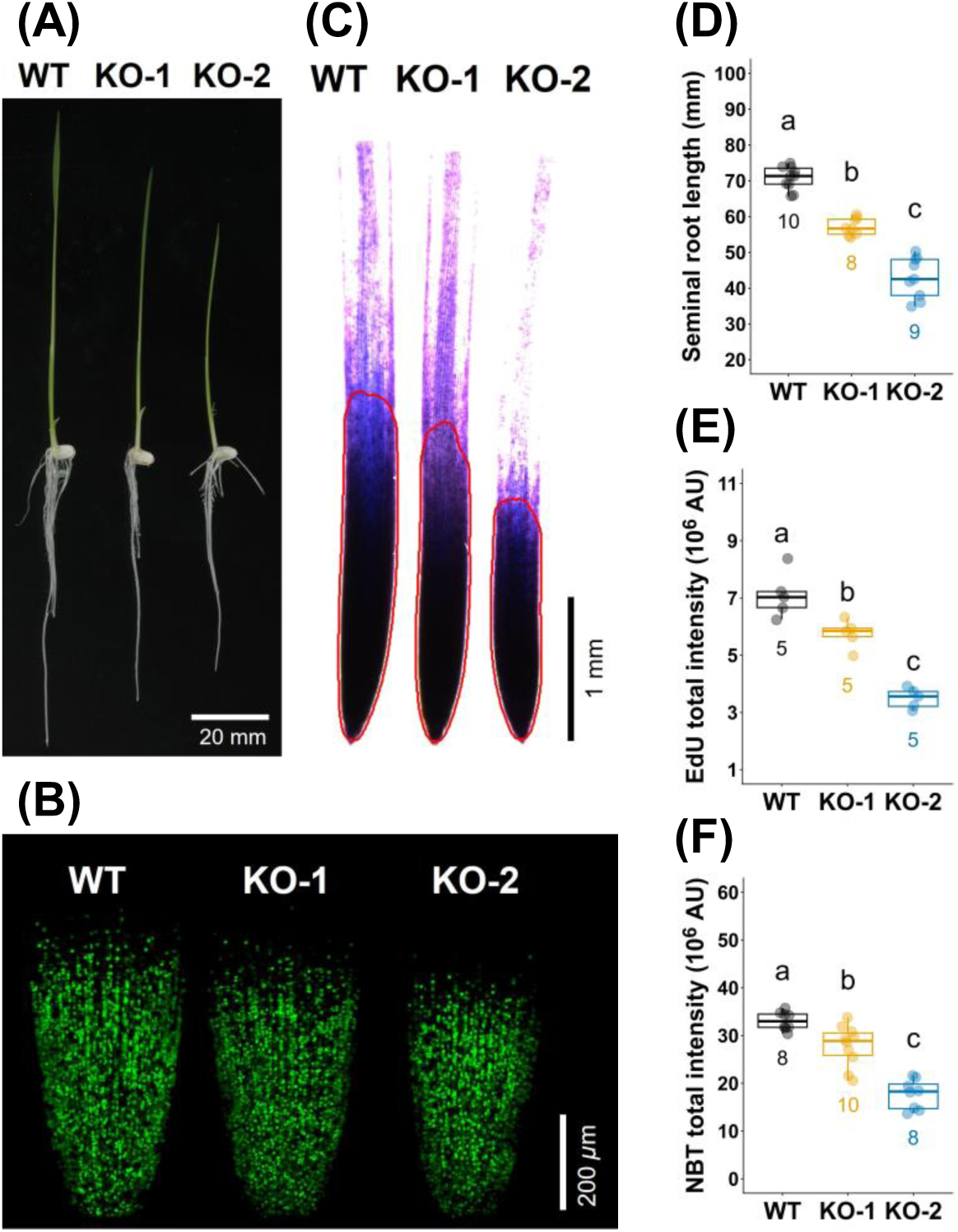
*Osrgf1-1* mutants exhibit shorter seminal roots, reduced root meristem activity, and reduced O₂•⁻ accumulation. (A) Representative photographs of 5-day-old wild-type (WT), *Osrgf1-1-7* (KO-1), and *Osrgf1-1-8* (KO-2) seedlings. Bar = 20 mm. (B) EdU-stained root meristem of WT, KO-1, and KO-2. Bar = 200 µm. (C) NBT-stained root tips of WT, KO-1, and KO-2. Red outlines indicate the NBT-stained root tip region used for quantification. Bar = 1 mm. (D) Seminal root length of WT, KO-1, and KO-2 seedlings. (E) EdU-stained total intensity in the root tips of WT, KO-1, and KO-2. (F) NBT-stained total intensity in the root tips of WT, KO-1, and KO-2. In (D), (E), and (F), box plots show the median, interquartile range, and individual data points. Numbers below indicate sample size (n). Different letters indicate statistically significant differences (Tukey’s HSD test, *P* < 0.05).

In *Arabidopsis*, reactive oxygen species (ROS) act downstream of RGF1 receptor signalling, and O₂•⁻ accumulates predominantly within the root meristem. To determine whether loss of OsRGF1-1 affects O₂•⁻ accumulation, roots were stained with nitroblue tetrazolium (NBT). Wild-type roots displayed broad and intense staining, whereas both mutants exhibited weaker and more restricted NBT signals (Fig. 3C, F), indicating reduced O₂•⁻ accumulation. Together, these results demonstrate that OsRGF1-1 is required for root meristem activity and O₂•⁻ accumulation in rice roots.

### The predicted mature OsRGF1-1 peptide is biologically active in rice roots and dose-dependently restores seminal root length and O₂•⁻ accumulation in the *Osrgf1-1* mutant

In *Arabidopsis*, AtRGF1 treatment increases both root meristem size and O₂•⁻ accumulation in a dose-dependent manner (Hsiao *et al*., 2025; Yamada *et al*., 2020); however, the biological activity of OsRGF1-1 in rice roots had not been experimentally examined. The *Osrgf1-1* mutant exhibited shorter seminal roots and reduced O₂•⁻ accumulation, presumably due to a lack of functional OsRGF1-1 peptide (Fig. 3). To determine the effective concentration of OsRGF1-1 required to restore these defects, *Osrgf1-1-8* mutant seedlings were treated with synthetic OsRGF1-1 peptide at 1, 10, 100, or 1000 pM, or mock solution, and seminal root lengths and O₂•⁻ accumulation were examined (Fig. 4). Seminal root lengths of seedlings treated with 1 and 10 pM OsRGF1-1 were comparable to those of mock-treated seedlings, indicating that these concentrations were insufficient to restore root growth (Fig. 4A, C). Treatment with 100 pM OsRGF1-1 significantly restored seminal root length to levels comparable to those of the wild type (Fig. 4A, C). In contrast, 1000 pM OsRGF1-1 reduced seminal root length below that of mock-treated mutants, suggesting an inhibitory effect at supraoptimal concentrations. Consistent with these root length data, O₂•⁻ accumulation was not significantly increased at 1 and 10 pM OsRGF1-1 (Fig. 4B, D). Treatment with 100 pM OsRGF1-1 significantly restored O₂•⁻ accumulation to levels comparable to those of the wild type (Fig. 4B, D). By contrast, 1000 pM OsRGF1-1 strongly elevated O₂•⁻ accumulation beyond wild-type levels (Fig. 4B, D), consistent with the inhibitory effect on root growth observed at this concentration.

**Fig. 4.**
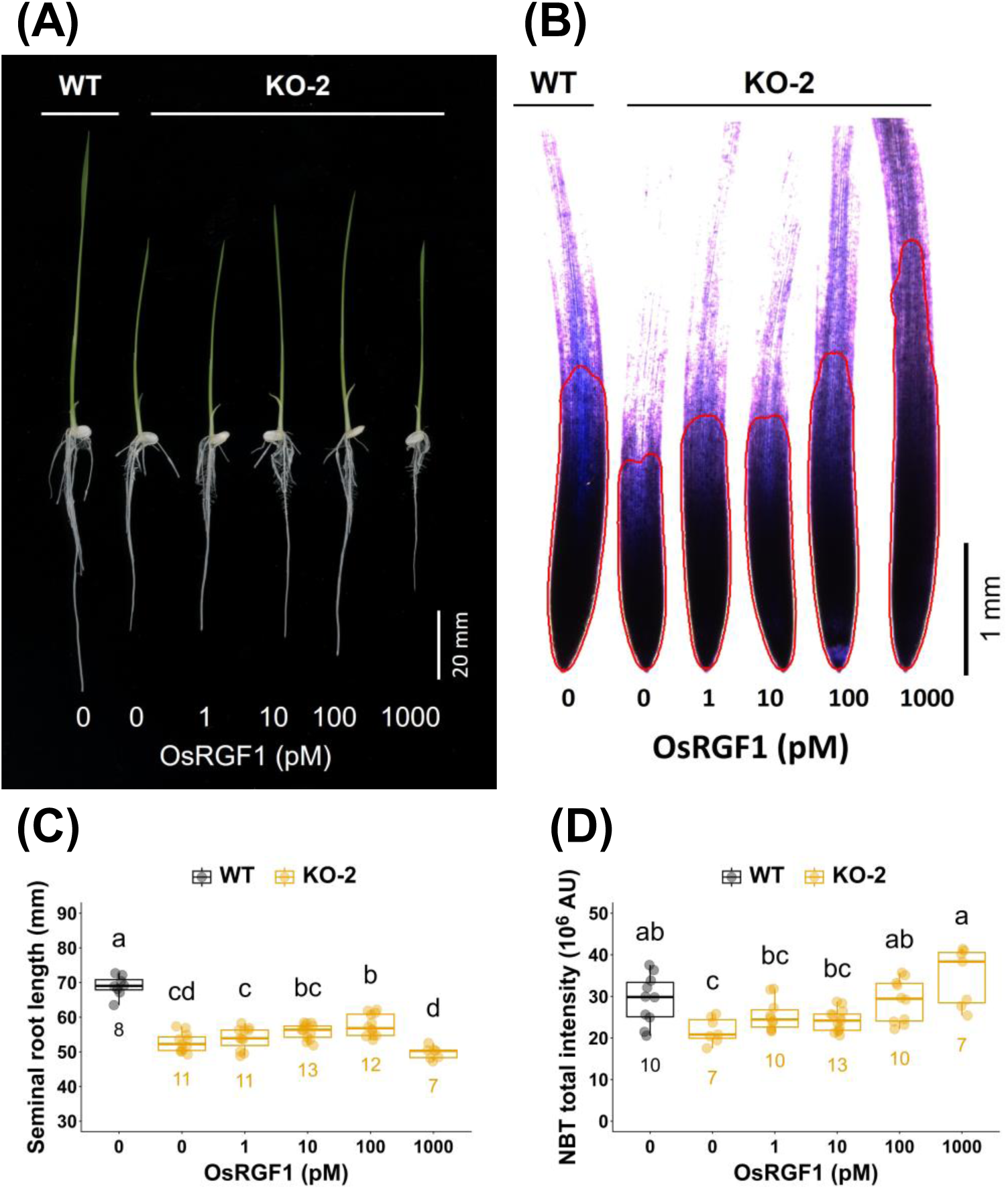
The *Osrgf1-1* mutant was sensitive to lower concentration of OsRGF1 peptide. (A) Representative photographs of 5-day-old wild-type (WT) and *Osrgf1-1-8* (KO-2) seedlings treated with 0, 1, 10, 100, or 1000 pM chemically synthesized OsRGF1-1 peptide. Bar = 20 mm. (B) NBT-stained root tips of seedlings treated as in (A). Red outlines indicate the NBT-stained root tip region used for quantification. Bar = 1 mm. (C) Primary root length of seedlings treated with increasing concentrations of OsRGF1-1 peptide. (D) NBT total intensity in root tips of seedlings treated as in (B). Box plots show median, interquartile range, and individual data points. Numbers below indicate sample size (*n*). Different letters indicate statistically significant differences (Tukey’s HSD test, *P* < 0.05).

To determine whether these concentrations of OsRGF1-1 affect wild-type roots, wild-type seedlings were treated with the same range of OsRGF1-1 concentrations (Fig. S2). Concentrations of 1, 10, and 100 pM OsRGF1-1 did not significantly affect seminal root length or O₂•⁻ accumulation in the wild type compared with mock treatment (Fig. S2), suggesting that endogenous OsRGF1-1 levels in wild-type roots are sufficient and that exogenous peptide at these concentrations does not exceed the optimal range. In contrast, 1000 pM OsRGF1-1 reduced seminal root length and elevated O₂•⁻ accumulation in both wild-type and *Osrgf1-1-8* mutant roots (Figs. 4, S2), indicating that this concentration is supraoptimal regardless of genotype. Notably, 100 pM OsRGF1-1 restored the *Osrgf1-1* mutant phenotypes but had no detectable effect on the wild type (Figs. 4, S2). This differential response strongly suggests that the *Osrgf1-1* mutant lacks sufficient endogenous OsRGF1-1 peptide, rendering it responsive to exogenous supplementation at concentrations that are below the threshold required to alter wild-type root development. Together, these results demonstrate that OsRGF1-1 exhibits a dose-dependent activity window, in which 100 pM is sufficient to restore seminal root growth and O₂•⁻ accumulation in the *Osrgf1-1* mutant, while supraoptimal concentrations exert inhibitory effects on both genotypes. These findings confirm that the mature OsRGF1-1 peptide is biologically active in rice roots, consistent with the known activity of AtRGF1 in *Arabidopsis*.

### OsRGF1-1 peptide treatment restores root defects in both alleles of *Osrgf1-1* mutants

To confirm that OsRGF1-1 is responsible for the root defects observed in *Osrgf1-1* mutants, we examined whether exogenous application of a chemically synthesised, sulfated OsRGF1-1 peptide could restore the mutant phenotypes in both alleles. As shown in Fig. 4, treatment with 100 pM OsRGF1-1 was the optimal concentration for rescuing the *Osrgf1-1-8* allele. We therefore examined whether this concentration similarly restores the root defects of both *Osrgf1-1* alleles (Fig. 5). Wild-type, *Osrgf1-1-7* (KO-1), and *Osrgf1-1-8* (KO-2) seedlings were treated with mock solution or 100 pM OsRGF1-1 peptide, and seminal root length and O₂•⁻ accumulation were examined (Fig. 5). Treatment with 100 pM OsRGF1-1 significantly promoted seminal root growth in both *Osrgf1-1* alleles compared with mock-treated controls (Fig. 5A, C). In contrast, seminal root growth in wild-type seedlings was unaffected by 100 pM OsRGF1-1 treatment (Fig. 5A, C). Similarly, 100 pM OsRGF1-1 increased O₂•⁻ accumulation in both *Osrgf1-1* alleles but had no detectable effect on the wild type (Fig. 5B, D). These results demonstrate that the root growth and O₂•⁻ accumulation defects observed in both CRISPR-generated alleles result from loss of functional OsRGF1-1 peptide, rather than off-target mutations.

**Fig. 5.**
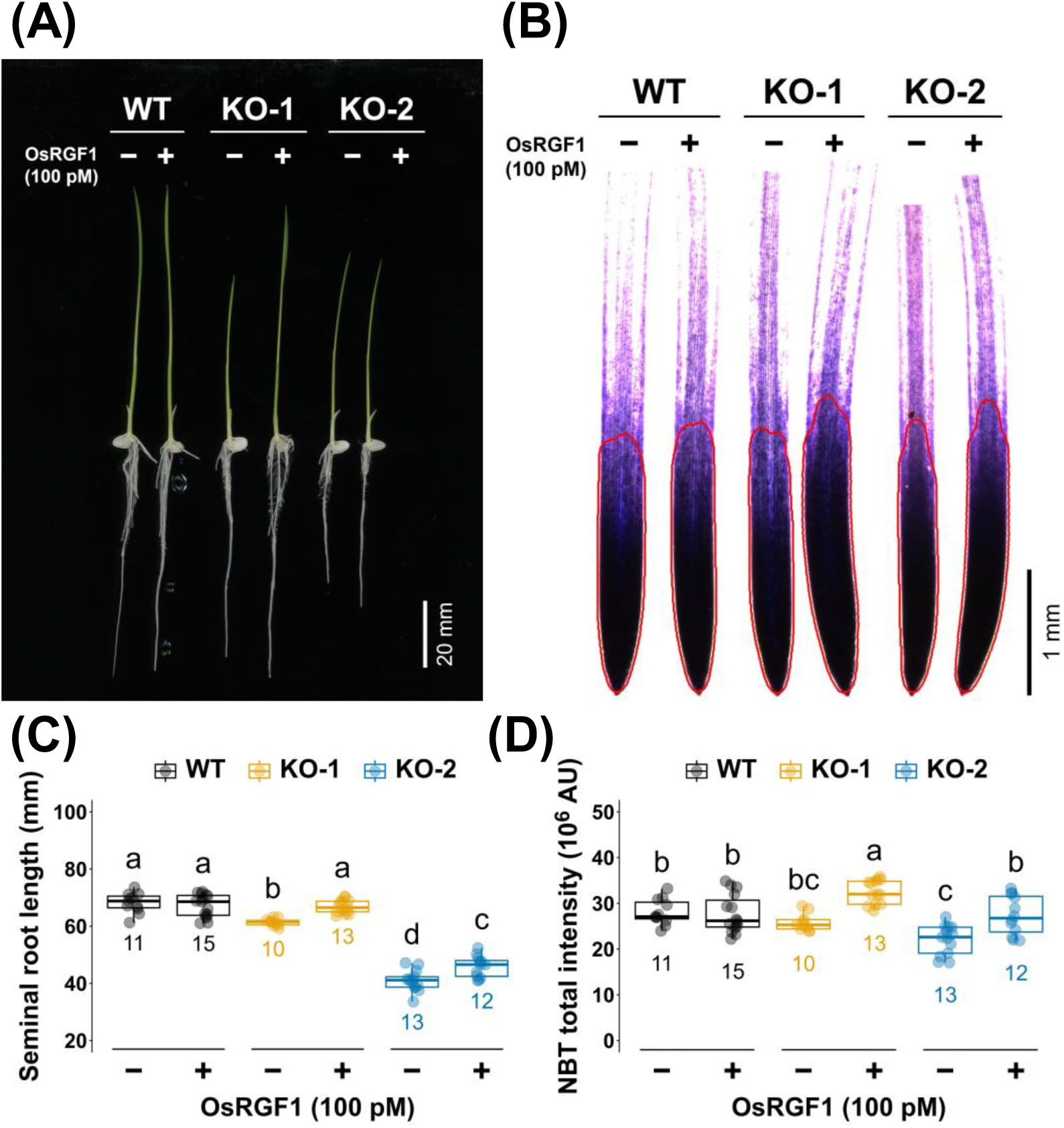
Exogenous OsRGF1-1 peptide rescues root growth and O₂•⁻ accumulation defects in *Osrgf1-1* mutants. (A) Representative photographs of 5-day-old WT, *Osrgf1-1-7* (KO-1), and *Osrgf1-1-8* (KO-2) seedlings treated with or without 100 pM OsRGF1-1 peptide. Bar = 20 mm. (B) NBT-stained root tips of seedlings treated as in (A). Red outlines indicate the NBT-stained root tip region used for quantification. Bar = 1 mm. (C) Primary root length of WT, KO-1, and KO-2 seedlings with or without OsRGF1-1 peptide treatment. (D) NBT total intensity in root tips of seedlings treated as in (B). Box plots show median, interquartile range, and individual data points. Numbers below indicate sample size (*n*). Different letters indicate statistically significant differences (Tukey’s HSD test, *P* < 0.05).

### OsRGF1-1 requires RGF receptors to enlarge root meristem size through ROS signaling

Since secreted RGF peptides require plasma membrane-localized receptors for signal transduction, we examined whether OsRGF1-1 signaling depends on known RGF receptors. Previous comparative genomic analyses have suggested the presence of AtRGF1 receptor homologues in the rice genome (Fang *et al*., 2021). To test whether OsRGF1-1 functions through the conserved RGF receptor pathway, we analyzed the responses of *Arabidopsis* wild-type and *rgfr1/2/3* mutant seedlings to OsRGF1-1 and AtRGF1 peptides.

In wild-type roots, treatment with 5 nM OsRGF1-1 or AtRGF1 similarly increased root meristem size (Fig. 6A, B) and enhanced O₂•⁻ accumulation, as detected by NBT staining (Fig. 6C, D). In contrast, neither OsRGF1-1 nor AtRGF1 treatment affected meristem size or O₂•⁻ accumulation in the *rgfr1/2/3* mutant (Fig. 6). In addition, the *rgfr1/2/3* mutant showed lower basal O₂•⁻ levels compared with the wild type. These results indicate that OsRGF1-1 requires RGF receptors to regulate O₂•⁻ accumulation and root meristem development.

**Fig. 6.**
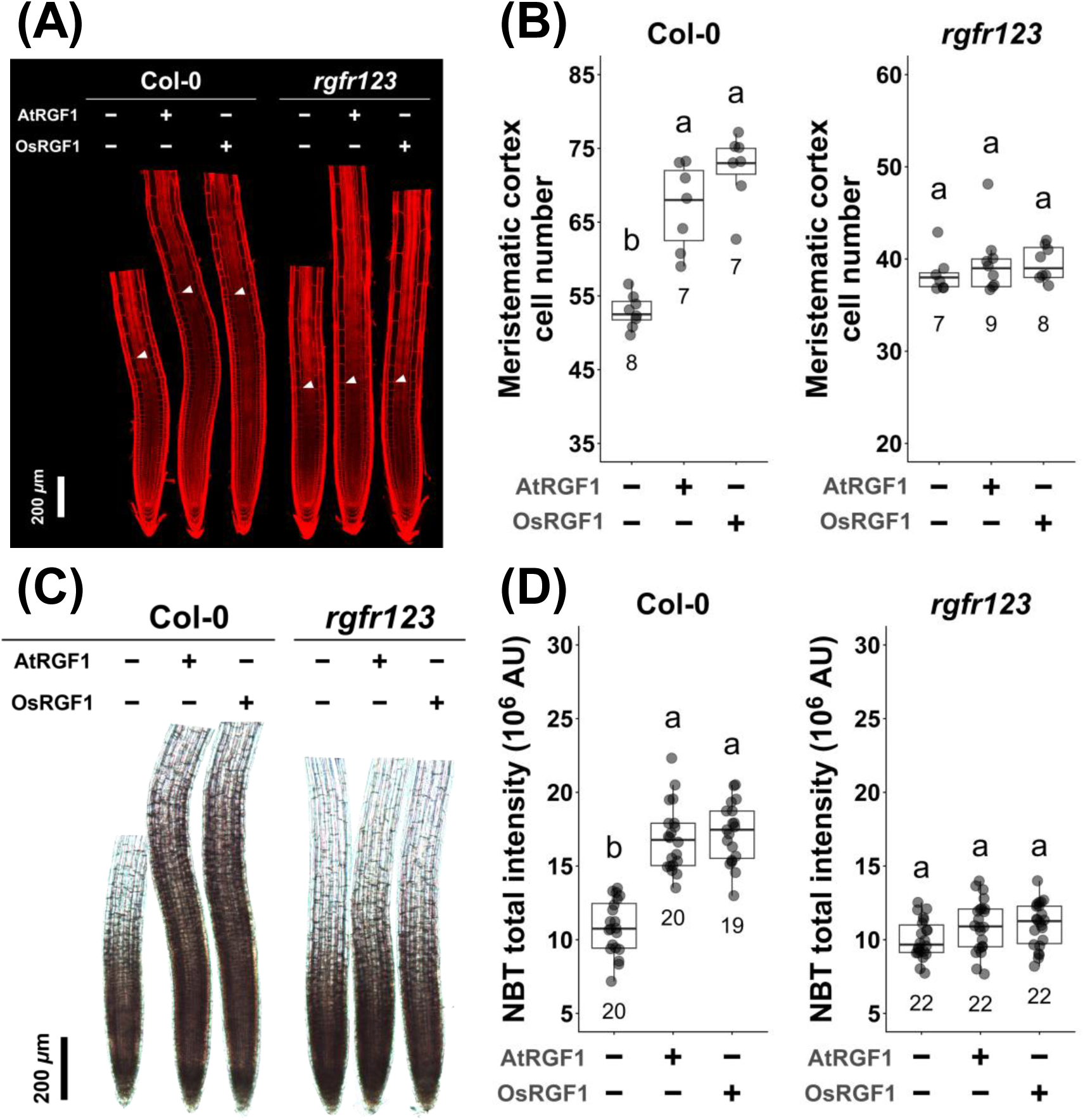
OsRGF1-1 signals through conserved RGF receptors in *Arabidopsis*. (A) Confocal images of propidium iodide-stained root tips of Col-0 and *rgfr123* seedlings treated with AtRGF1 or OsRGF1-1 peptide (5 nM). Arrowheads indicate the boundary of the meristematic zone. Bar = 200 µm (B) Meristematic cortex cell number in Col-0 and *rgfr123* roots treated as in (A). (C) NBT-stained root tips of Col-0 and *rgfr123* seedlings treated with AtRGF1 or OsRGF1-1 peptide. Bar = 200 µm (D) Total NBT intensity in root tips of Col-0 and *rgfr123* seedlings treated as in (C). In (B) and (D), box plots show median, interquartile range, and individual data points. Numbers below indicate sample size (*n*). Different letters indicate statistically significant differences (Tukey’s HSD test, *P* < 0.05).

Together, these results demonstrate that OsRGF1-1 functions as a positive regulator of root meristem development through RGF receptor-dependent ROS signaling.

## Discussion

In *Arabidopsis*, RGF peptides regulate root meristem size by controlling ROS levels through receptor-mediated signalling. Our findings extend this framework to rice, showing that OsRGF1-1 functions through a conserved peptide–ROS signalling mechanism linking peptide perception to O₂•⁻-dependent redox control of the root meristem. In this context, OsRGF1-1 appears to function as part of a conserved regulatory mechanism that integrates extracellular peptide signals with intracellular redox states to maintain meristem homeostasis.

### OsRGF1-1 represents a functional homologue of AtRGF1

Previous phylogenetic analyses have identified four homologues of AtRGF1 in the *Oryza sativa* genome (Ghorbani *et al*., 2015; Fang *et al*., 2021). Among these, OsRGF1-1 has been repeatedly identified as the closest rice homologue of AtRGF1 and shares key features with AtRGF1, including conserved DY, HPP, and N residues, a C-terminally localised mature peptide, and a predicted 13-amino acid active sequence (Fig. 1). By contrast, the other previously predicted homologues show limited conservation of characteristic AtRGF1 features, including mature peptide length and conserved residues. One possible explanation is that earlier studies relied on older genome annotations, which may have affected homologue prediction accuracy. It also remains possible that future rice genome annotations and comparative genomic analyses will identify additional RGF-related peptides. In *Arabidopsis*, AtRGF genes exhibit distinct expression domains associated with their roles in root development, with *AtRGF1/2/3* redundantly expressed around the root apical meristem (Fernandez *et al*., 2013). Similarly, OsRGF1-1 displayed a highly restricted expression pattern near the root apex of rice seminal and crown roots (Fig. 1B,C, and D), consistent with publicly available single-cell RNA-seq data showing preferential expression in meristematic cell populations (Fig. 1 B,C, and D). Loss of OsRGF1-1 function leads to defects in seminal root growth, meristem size, and O₂•⁻ accumulation (Fig. 3).

Together, these phylogenetic, structural, expression, and functional data support the conclusion that OsRGF1-1 acts as a functional homologue of AtRGF1 in regulating root meristem development in rice.

The apparent central role of OsRGF1-1 in rice raises the question of why a single RGF1-like gene plays a major role, whereas multiple RGF peptides act redundantly in *Arabidopsis*. One possible explanation is that the RGF family has undergone distinct evolutionary trajectories in monocots and dicots, resulting in differences in gene retention, subfunctionalisation, or expression partitioning. In rice, the restricted expression of OsRGF1-1 within the root meristem may confer functional specificity and limit redundancy with other predicted RGF homologues. Alternatively, other RGF-like peptides may function in different developmental contexts or under specific environmental conditions.

Consistent with this idea, although the *Osrgf1-1* single mutant exhibits clear defects in seminal root growth and meristem size, the phenotype is less severe than that observed in higher-order *rgf* mutants in *Arabidopsis*. This suggests that additional RGF-like peptides may contribute to meristem regulation in rice, potentially acting in partially redundant or parallel pathways. Future transcriptome analyses of the *Osrgf1-1* mutant may help identify such components by revealing compensatory or parallel signalling responses.

Although both *Osrgf1-1* alleles showed significant responses to exogenous OsRGF1-1 peptide treatment, the *Osrgf1-1-8* allele exhibited a more severe root phenotype than *Osrgf1-1-7* (Figs. 3, 5). Both alleles are predicted to lack the functional mature OsRGF1-1 peptide; however, unlike *Osrgf1-1-7*, which carries a premature stop codon upstream of the mature peptide region, *Osrgf1-1-8* is predicted to produce a truncated precursor polypeptide. It is possible that this aberrant polypeptide may interfere with root development in a manner distinct from simple loss of function, contributing to the more severe phenotype observed in *Osrgf1-1-8*. Nevertheless, since both alleles were rescued by exogenous OsRGF1-1 peptide application, the primary cause of the root defects is loss of functional OsRGF1-1 signalling.

### Functional evidence supports OsRGF1-1 as an orthologue of AtRGF1

The biological activity of the mature 13-amino acid OsRGF1-1 peptide, together with its functional interchangeability with AtRGF1 in *Arabidopsis*, supports the conclusion that OsRGF1-1 is perceived through a conserved receptor-mediated signalling pathway (Fig. 6). The inability of both OsRGF1-1 and AtRGF1 to elicit responses in *Arabidopsis* RGF receptor mutants (Fig. 6) indicates that OsRGF1-1 is recognised by the same receptor machinery as AtRGF1. Moreover, chemical complementation of *Osrgf1-1* mutants by 100 pM OsRGF1-1 (Fig. 5) demonstrates that the processed peptide is biologically active *in vivo*. Notably, 100 pM OsRGF1-1 was sufficient to rescue the *Osrgf1-1* mutant phenotypes, whereas 1000 pM was required to elicit a detectable response in wild-type roots. This difference is consistent with the idea that *Osrgf1-1* mutants have reduced endogenous peptide levels, rendering them more sensitive to exogenous OsRGF1-1 supplementation. Taken together, these findings support the conclusion that OsRGF1-1 functions as a rice orthologue of AtRGF1, regulating root meristem size through receptor-dependent ROS signalling. Furthermore, the ability of the sulfated OsRGF1-1 peptide to restore *Osrgf1-1* mutant phenotypes in rice suggests that key aspects of peptide processing and post-translational modification are conserved between dicots and monocots. In *Arabidopsis*, the molecular basis of this ROS-dependent regulation has recently been elucidated: H₂O₂ induces S-sulfenylation of the 212th cysteine residue of PLT2, leading to its degradation, while O₂•⁻ in the meristematic zone maintains PLT2 stability by preserving the free thiol state of this residue (Hsiao *et al*., 2025). Whether a similar cysteine-dependent redox regulation of PLT2 homologues operates in rice represents an important question for future investigation.

Together, our findings define OsRGF1-1 as a peptide signal that links extracellular peptide perception to intracellular redox control of root meristem activity in rice. By integrating genetic and cross-species functional evidence, this study establishes OsRGF1-1 as a functional rice orthologue of AtRGF1 within an evolutionarily conserved peptide–receptor–ROS signalling module. Furthermore, recent work has demonstrated that the RGF–receptor–PLT2 pathway plays a central role in root adaptation to non-lethal thermal stress in *Arabidopsis*, operating independently of the canonical heat shock response (Hsiao *et al*., 2026). Given that OsRGF1-1 functions as a conserved rice homologue of AtRGF1, it will be of considerable interest to determine whether OsRGF1-1 similarly mediates root meristem adaptation to heat stress in rice, a crop of major agricultural importance. At the same time, the distinct genetic organisation and deployment of RGF peptides in rice and *Arabidopsis* suggest that conserved signalling mechanisms have been adaptively rewired during monocot and dicot evolution.

## Supplemental figures

**Fig. S1.**
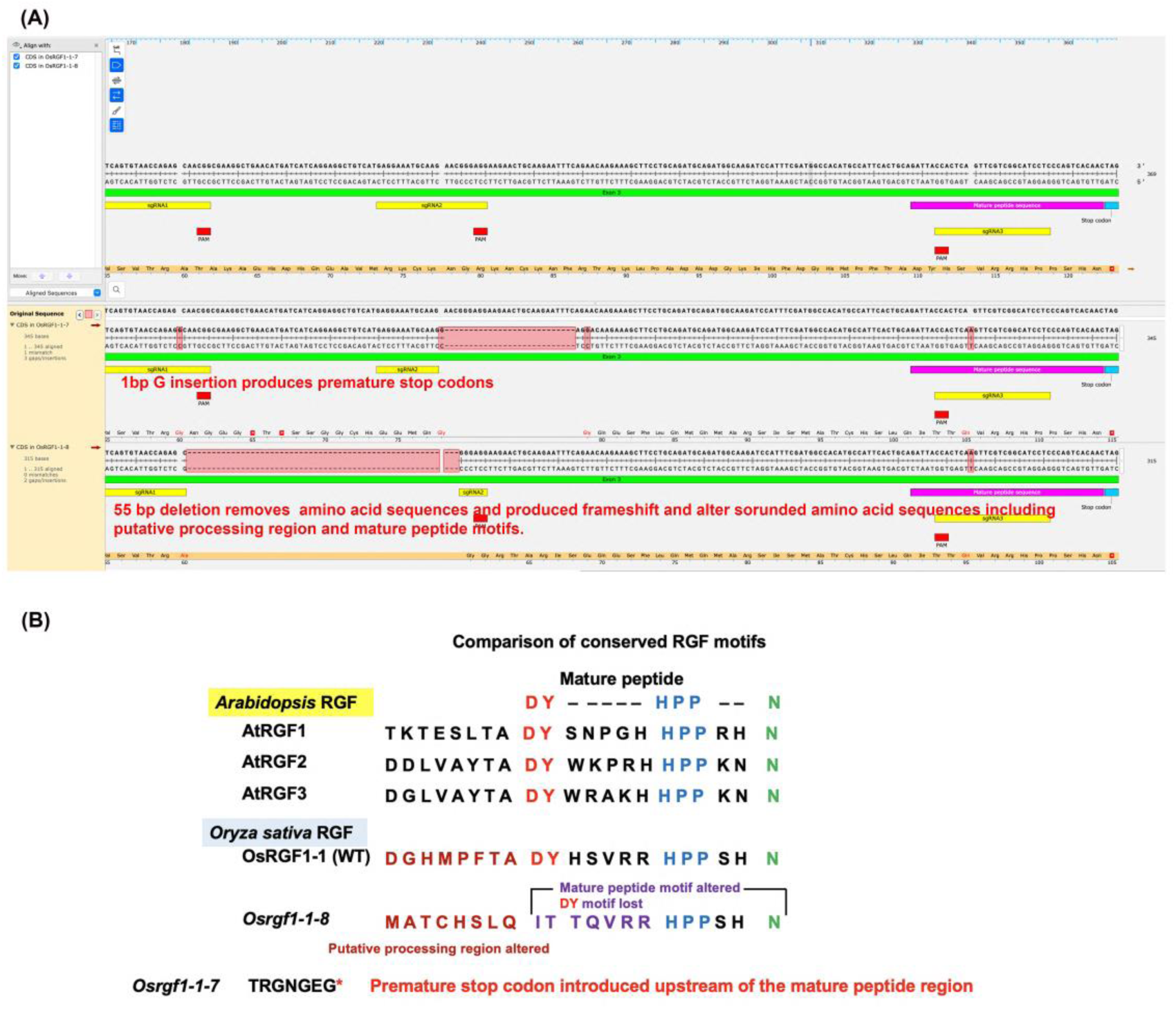
Characterisation of CRISPR/Cas9-generated *Osrgf1-1* mutant alleles. (A) Schematic of the *OsRGF1-1* genomic sequence showing the positions of three guide RNAs (sgRNA1, sgRNA2, sgRNA3) and the PAM sequences. The mature peptide sequence is indicated. Mutations identified in the *Osrgf1-1-7* and *Osrgf1-1-8* alleles are shown with red highlighting. The *Osrgf1-1-7* allele carries a 1-bp G insertion producing premature stop codons. The *Osrgf1-1-8* allele carries a 55-bp deletion that removes amino acid sequences and produces a frameshift, altering surrounding amino acid sequences including the putative processing region and mature peptide motifs. (B) Alignment of the predicted mature peptide sequences of *Arabidopsis* RGF1, RGF2, and RGF3 with OsRGF1-1 (WT) and the *Osrgf1-1* mutant alleles. Conserved DY (red) and HPP (blue) motifs and the terminal asparagine N (green) are indicated. In *Osrgf1-1-8*, the mature peptide motif is altered and the DY motif is lost. In *Osrgf1-1-7*, a premature stop codon is introduced upstream of the mature peptide region.

**Fig. S2.**
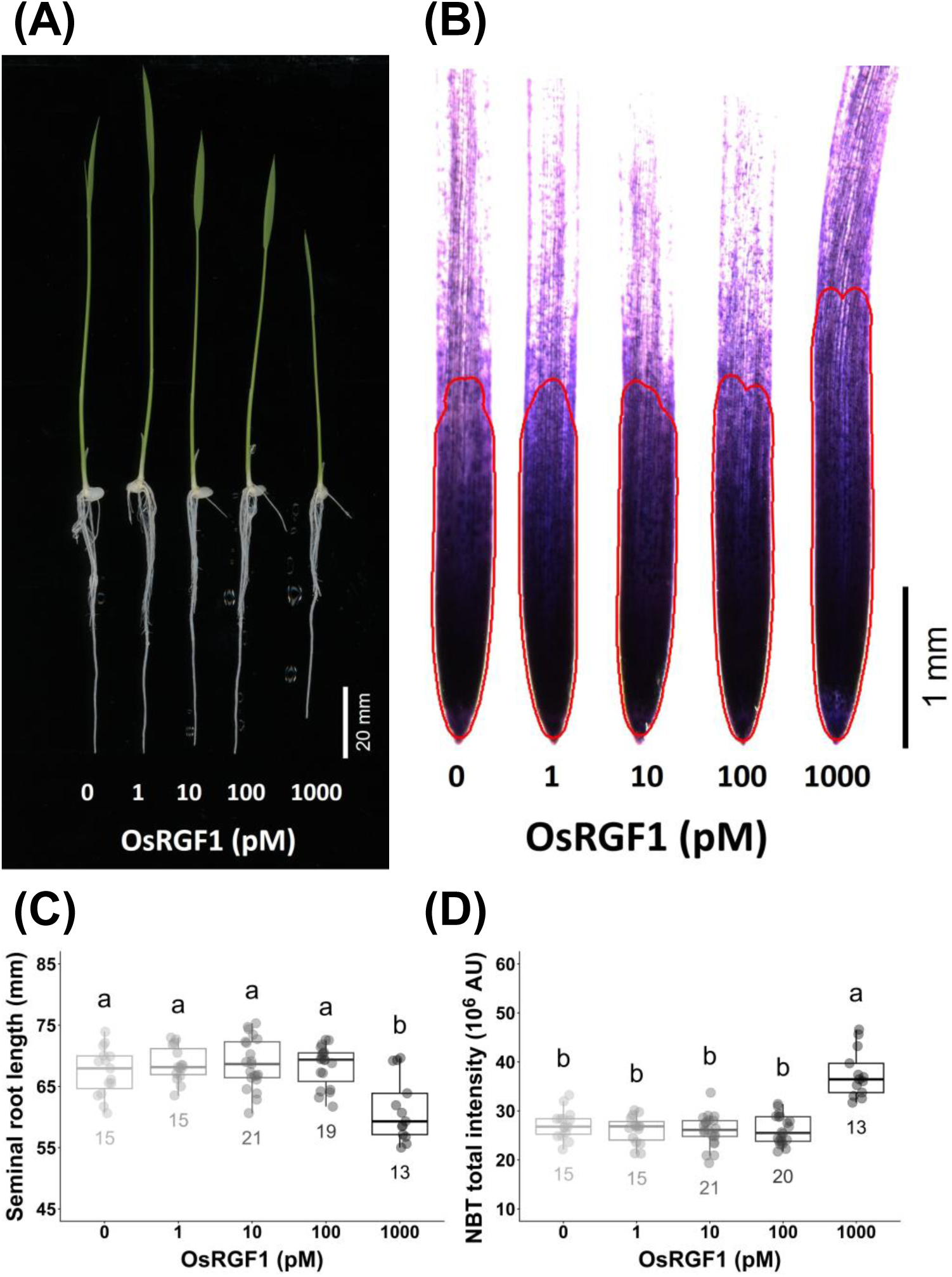
Exogenous OsRGF1-1 peptide promoted O₂•⁻ accumulation in a dose-dependent manner. (A) Representative photographs of 5-day-old wild-type (WT) seedlings treated with 0, 1, 10, 100, or 1000 pM chemically synthesized OsRGF1-1 peptide. Bar = 20 mm. (B) NBT-stained root tips of seedlings treated as in (A). Red outlines indicate the NBT-stained root tip region used for quantification. Bar = 1 mm. (C) Primary root length of seedlings treated with increasing concentrations of OsRGF1-1 peptide. (D) NBT total intensity in root tips of seedlings treated as in (B). Box plots show median, interquartile range, and individual data points. Numbers below indicate sample size (*n*). Different letters indicate statistically significant differences (Tukey’s HSD test, *P* < 0.05).

## Acknowledgements

We thank the Yamada lab members and Jian-You Wang for valuable comments on the manuscript; Fu-Hui Wu and Choun-Sea Lin at the Plant Tech. Core Laboratory of Agricultural Biotechnology Research Center (ABRC), Academia Sinica for generation of CRISPR/Cas9 construct; Lin-Yun Kuang at ABRC, Academia Sinica for transformation the CRISPR/Cas9 construct into rice; the Confocal Microscope Core Facility at the Biotechnology Center in Southern Taiwan, Academia Sinica, for maintaining the confocal laser scanning microscope.

This work was supported by a Career Development Award, Academia Sinica, Taiwan (AS-CDA-111-L05); the National Science and Technology Council, Taiwan (112-2311-B-001-021-MY3, 111-2311-B-001-028, 109-2313-B-001-006-MY3); Innovative Translational Agricultural Research Program, Academia Sinica (AS-KPQ-110-ITAR-13), and the Agricultural Biotechnology Research Center, Academia Sinica, Taiwan, to M.Y.

## Author contributions

M.Y. and J-K. L. conceptualized the study; J-K. L., J-N. J., H-C. Y., H-Y. C., Y-C.H., H. B., and C-S. T. performed all experiments; J-K. L. performed the statistical analyses; all the authors wrote the paper.

